# Nocturnal increase in cerebrospinal fluid secretion as a regulator of intracranial pressure

**DOI:** 10.1101/2023.03.14.532581

**Authors:** Annette Buur Steffensen, Beatriche Louise Edelbo, Dagne Barbuskaite, Søren Norge Andreassen, Markus Harboe Olsen, Kirsten Møller, Nanna MacAulay

## Abstract

It is crucial to maintain the intracranial pressure (ICP) within the physiological range to ensure proper brain function. The ICP may fluctuate during the light-dark phase cycle, complicating diagnosis and treatment choice in patients with pressure-related disorders. Such ICP fluctuations may originate in circadian or sleep-wake cycle-mediated modulation of cerebrospinal fluid (CSF) flow dynamics, which in addition could support diurnal regulation of brain waste clearance. Through a combination of patient data and *in vivo* telemetric pressure measurements in adult male rats, we demonstrated that ICP increases in the dark phase in both species, independently of vascular parameters. This increase aligns with elevated CSF collection in patients and CSF production rate in rats, the latter obtained with the ventriculo-cisternal perfusion assay. The dark-phase increase in CSF secretion in rats was, in part, assigned to increased transport activity of the choroid plexus Na^+^,K^+^,2Cl^-^ cotransporter (NKCC1), which is implicated in CSF secretion by this tissue. In conclusion, CSF secretion, and thus ICP, increases in the dark phase in humans and rats, irrespective of their diurnal/nocturnal activity preference, in part due to altered choroid plexus transport activity. Our findings suggest that CSF dynamics are modulated by the circadian rhythm, rather than merely sleep itself.

## BACKGROUND

Our skull represents a fixed-volume compartment enclosing the brain tissue and the fluids that surround its cells and structures; the cerebrospinal and interstitial fluids (CSF and ISF) as well as the blood that supplies the brain with oxygen and nutrients. The Kellie-Monroe doctrine stipulates that due to the constant intracranial volume, the volumes of its components are in a dynamic equilibrium, meaning that an increase in one component is normally compensated by a decrease in another, in order to ensure a stable intracranial pressure (ICP) [1,2]. When these compensatory mechanisms are exhausted, the ICP will increase if the intracranial volume increases further, e.g. during expansion of a hematoma, progressive hydrocephalus, or cerebral oedema. The brain is vulnerable to increased ICP both because of the risk of herniation, i.e., mechanically induced injury as tissue becomes squeezed against naturally occurring anatomical borders, and because it reduces cerebral perfusion pressure, potentially leading to ischemia [3]. Accordingly, the clinical care of severe acute brain injury, as well as other conditions featuring intracranial volume disturbances, regularly involves ICP monitoring and may include CSF drainage to manage the elevated pressure. The ICP appears to fluctuate with the time of day in human patients with a nightly elevation of this parameter [4–6], which may influence diagnosis and treatment of pressure-related disorders. However, with the ICP fluctuations occurring with posture changes [7,8], such nightly ICP elevation may, in part, occur with the transition from vertical to horizontal position associated with sleep. It therefore remains unresolved i) to what extent the ICP inherently fluctuates with the time of day, and if so, ii) whether the ICP fluctuations arise as a consequence of altered CSF dynamics, and iii) whether such fluctuations reside with the sleep-wake cycle or are dictated by the circadian rhythm.

Sleep increases the interstitial space in rodent parenchyma and is permissive for cortical CSF tracer influx [9]. The sleep-induced increase in CSF tracer penetration in the brain is predicted to facilitate removal of cellular waste products and toxins [9,10]. This process is thought enhanced by the increased low-frequency pulsation in CSF flow observed during non-rapid eye movement (NREM) sleep in humans [11]. An increased rate of CSF secretion during sleep could support the proposed metabolic waste clearance. Accordingly, magnetic resonance imaging (MRI) of the pulsatile flow of CSF through the cerebral aqueduct in humans suggested a nightly increase in the net rate of CSF flow as a proxy for CSF secretion [12]. However, a subsequent study employing a modified MRI approach failed to observe such nightly elevation in CSF flow [13], rendering the possibility of a diurnally fluctuating CSF secretion rate unresolved.

Thus, we set out to quantify day-night ICP fluctuations and resolve the underlying CSF dynamics in humans and experimental rats. Our results reveal that CSF secretion increases in the dark phase, which may contribute to the elevated ICP observed in this phase. The dark phase increase in CSF dynamics occurred in both species, irrespective of their diurnal/nocturnal activity preference, and our findings thus indicate that CSF dynamics might not be sleep dependent as such, but rather modulated by the circadian rhythm.

## METHODS

### Patients

For the present study, patients who were admitted between 1 August, 2017 and 31 December, 2020 and who underwent placement of an external ventricular drain (EVD) and continuous measurement of ICP, could be included. From this group, 50 patient data sets were retrospectively and randomly selected from a patient group with various diagnoses of acute brain injury. Of these 50 data sets, 25 fulfilled the study criteria that i) ICP measurements were available during both night and day in the same patient; ii) CSF was drained against an unchanged counter-pressure (drain height) for at least four consecutive days, iii) CSF output measurements were stable (defined by less than 10% daily variation in the CSF collection). One of these patients was subsequently excluded since one ICP measurement was above 100 mmHg, which was considered out of normal patient range and potentially could reflect a detection failure. The patient data set employed for the analysis originated from 24 patients (median age: 53; range 21–83, 13F/11M) with the following diagnoses: non-traumatic subarachnoid haemorrhage (n = 12), spontaneous intracerebral haemorrhage (n=6), and traumatic brain injury (n=6), who were mainly bedridden during the time of measurements, either in the supine position or with a 30–45° headrest elevation. At admission, the patients scored 9.0 ± 4.5 on the Glascow Coma scale [14] and patient monitoring commenced 4.3 ± 3.5 days after ictus. Clinical management followed international guidelines [15–17]. The ICP was measured either with a Codman intraparenchymal sensor (Integra LifeSciences, Princeton, New Jersey, USA) or a Spiegelberg combidrain (Spiegelberg GmbH & Co., Hamburg, Germany) with the zero reference point at the external auditory meatus, and the CSF flow estimated from the hourly fluid output from the EVD at a given drain resistance. Of 24 included patients, in 21 blood pressure was measured invasively through the radial artery with the zero reference point at the external auditory meatus to allow for calculation of (and therapy guided by) cerebral perfusion pressure. The remaining three patients had no continuous blood pressure measurements available during the periods of interest. The physiological variables were obtained for the light period (defined as 6 AM to 6 PM) and for the dark period (defined as 6 PM to 6 AM) and ‘fluctuations’ refer to the differences between night and day measurements.

### Experimental rats and anesthesia regimen

Male Sprague-Dawley rats 9–11 weeks old, housed in temperature-controlled facilities with 12:12 hour light cycle, were employed for the study. The rats had access to water and food *ad libitum* and were randomly assigned to experimental groups. For terminal procedures, rats were anaesthetized with 6 mg/ml xylazine + 60 mg/ml ketamine (ScanVet) in sterile water (0.17 ml/100 g body weight through intraperitoneal injection). When prolonged anesthesia was required (CSF secretion measurements), the animals received bolus ketamine injections at half the initial dose as required to sustain anesthesia. A tracheotomy was performed to control ventilation using the VentElite system (Harvard Apparatus) by 0.9 l/min humidified air mixed with 0.1 l/min O_2_ adjusted with approximately 3 ml per breath, 80 breath/min, a Positive End-Expiratory Pressure (PEEP) at 2 cm, and 10% sigh for a ∼400 g rat. Telemetric pressure probe insertion procedures were performed using isoflurane (Attane vet, 1000 mg/g isoflurane, ScanVet) mixed with 1.8 l/min air / 0.1 l/min O_2_. Body temperature during all surgeries were maintained at 37°C by a homoeothermic monitoring system (Harvard Apparatus).

### Telemetric pressure measurements in rats

Implantation was performed as described previously [18,19] using KAHA Sciences rat dual pressure telemetric system. Analgesia was injected s.c. prior and post-surgery (48 h) and comprised 5 mg/kg Caprofen (Norodyl Vet, Norbrook), 0.05 mg/kg buprenorphine (Tamgesic, Indivior), and 200 + 40 mg/kg sulfadiazin and trimethoprim (Borgal Vet), and the surgical areas sterilized using 0.5 % chlorhexidine (Medic). An abdominal midline incision was made to isolate the abdominal aorta for implantation of the probe for mean arterial pressure and heart rate measurements, which was secured in the aorta using tissue adhesive (Histoacryl, Enbucrilate; B. Braun) and surgical mesh (PETKM2002, SurgicalMesh; Textile Development Associates). The body of the telemetric device was fixed to the abdominal wall with 4–0 non-absorbable EthilonII suture (Ethicon). The second pressure probe was tunneled, using a 6 mm diameter stainless steel straw (Ecostrawz), to the base of the skull. After placing the animal in a stereotactic frame, the skull was exposed and two holes (1.2 mm) for anchor screws (00-96 × 3/32, Bilaney Consultants GmbH), as well as a 1.4 mm drill hole for the probe, were drilled posterior to bregma. The probe was placed epidurally in between the contralateral fastened screws, the probe hole was filled with spongostan (Ethicon). The probe was secured using surgical mesh and tissue adhesive, and lastly dental impression material (Take 1 Advanced, Kerr) was applied over the catheter and screws. The head incision was sutured using 4-0 non-absorbable EthilonII suture, the abdominal muscles secured using 4-0 absorbable Vicryl suture (Ethicon), and abdominal skin incision closed with suture clips (11 × 2 mm Michel clips, B. Braun). After waking, animals were placed in their cages on the TR181 Smart Pads (Kaha Sciences) and data were acquired at 1 kHz with PowerLab and LabChart software (v8.0, ADInstruments) when a stable baseline was achieved. Data presented are from baseline recordings obtained prior to initiation of a separate study [19]. To obtain times at which the animals were sedentary, the animals were filmed with a wireless camera (Cell Pro, Imou life) in the time intervals from 12-5 PM/AM. Within these intervals, time slots were chosen where the animals did not move for at least 30 s, and the ICP values determined in a blinded manner.

### CSF secretion rate measurements in rats

The CSF production rate was determined with the ventriculo-cisternal perfusion technique, with a total number of 12 animals used. One animal died during experimentation and was excluded alongside its day-night phase matching counterpart. An infusion cannula (Brain infusion kit 2, Alzet) was stereotaxically placed in the right lateral ventricle of an anesthetized and ventilated rat (1.3 mm posterior to Bregma, 1.8 mm lateral to the midline, and 0.6 mm ventral through the skull), through which pre-heated (37°C, SF-28, Warner Instruments) HCO_3_--containing artificial cerebrospinal fluid (aCSF; (in mM) 120 NaCl, 2.5 KCl, 2.5 CaCl_2_, 1.3 MgSO_4_, 1 NaH_2_PO_4_, 10 glucose, 25 NaHCO_3_, pH adjusted with 95% O_2_/5% CO_2_, 307 ± 1 mOsm) containing 1 mg/ml TRITC-dextran (tetramethylrhodamine isothiocyanate-dextran, MW = 155 kDa; T1287, Sigma), was perfused at 9 μl/min. CSF was sampled at 5 min intervals from cisterna magna with a glass capillary (30-0067, Harvard Apparatus pulled by a Brown Micropipette puller, Model P-97, Sutter Instruments) placed at a 5° angle (7.5 mm distal to the occipital bone and 1.5 mm lateral to the muscle-midline). The cisterna magna puncture and associated continuous fluid sampling prevents elevation of the ICP during the procedure. The dilution of the infused solution is ascribed to endogenously secreted CSF, irrespective of origin of this fluid [20]. The fluorescent content was measured in a microplate photometer (545 nm, Synery™ Neo2 Multi-mode Microplate Reader; BioTek Instruments), and the production rate of CSF was calculated from the equation [21]: V_p_ = r_i_ × (C_i_ – C_o_ / C_o_), where V_p_ = CSF production rate (μl/min), r_i_ = infusion rate (μl/min), C_i_ = fluorescence of inflow solution, and C_o_ = fluorescence of outflow solution. For the ventriculo-cisternal perfusion assay conducted during the dark phase, the anesthesia and all parts of the experiments were conducted either in the dark or with a red LED lamp for dark adaptive lighting (OcuScience), with the animals’ eyes completely covered following anesthesia, thus ensuring that the animals had minimal exposure to light. The CSF secretion rates were obtained approximately 8 hours after initiation of the light/dark phase (zeitgeber time (ZT) 8/20 h).

### RNAseq of rat choroid plexus

Rat choroid plexus (lateral and 4^th^) were isolated in the light phase (8 hours after light phase initiation, ZT 8 h) and in the dark phase (8 hours after dark phase initiation, ZT 20 h), the latter in the dark with only a red LED lamp for dark adaptive lighting (OcuScience) occasionally lit to ensure that light exposure was minimal during anesthesia and brain isolation. 12 animals were used for these experiments. The choroid plexuses were stored in RNAlater (Sigma) at -80 °C prior to RNA extraction and library preparation (performed by Novogene Company Limited with NEB Next® Ultra™ RNA Library Prep Kit (NEB)) prior to RNA sequencing (paired-end 150 bp, with 12 Gb output) on an Illumina NovaSeq 6000 (Illumina). Quality control and adapter removal were done by Novogene. The 150 base paired-end reads were mapped to rat reference genome Rnor_6.0.104 (*Rattus norvegicus*) using Spliced Transcripts Alignment to a Reference (STAR) RNA-seq aligner (v.2.7.9a) [22,23]. The mapped alignment generated by STAR was normalized to transcripts per million (TPM) with RSEM (v 1.3.3) [23,24]. Gene information was collected from ensembl’ BioMart [25]. The count table with TPM normalization were used for further analysis. Scripts and program parameters can be found at: https://github.com/Sorennorge/MacAulayLab_Nocturnal_rythem_RNAseq.

### Radioisotope flux assays

Rat lateral choroid plexuses were isolated in the light phase (8 hours after light phase initiation, ZT 8 h) and in the dark phase (8 hours after dark phase initiation, ZT 20 h), the latter in the dark with only a red LED lamp for dark adaptive lighting (OcuScience) occasionally lit during anesthesia and brain isolation to ensure that light exposure was minimal. The choroid plexus was placed in HEPES-aCSF (containing (in mM) 120 NaCl, 2.5 KCl, 2.5 CaCl_2_, 1.3 MgSO_4_, 1 NaH_2_PO_4_, 10 glucose, 17 Na-HEPES (4-(2-hydroxyethyl)-1-piperazineethanesulfonic acid, pH 7.56, 37°C) for 10 min prior to initiation of the experiment. ^86^Rb^+^ influx: The experiments were initiated by placing the choroid plexus in HEPES-aCSF containing 1 μCi/ml ^86^Rb^+^ (022-105721-00321-0001, POLATOM, as a tracer for K^+^ transport) and 4 μCi/ml ^3^H-mannitol (Perkin Elmer, as an extracellular marker [26]) at 37°C for 2 min in the absence or presence of 2 mM ouabain (O3125, Sigma). The two lateral choroid plexuses from each animal were randomly assigned to the control and ouabain conditions. The choroid plexus was subsequently swiftly rinsed in cold isotope-free HEPES-aCSF containing 2 mM ouabain (4°C) and transferred to a scintillation vial and dissolved in 100 μl Solvable (6NE9100, Perkin Elmer). For the light and dark phase influx experiment, 12 animals was used (six either phase), but one dark phase lateral choroid plexus (ouabain) was lost during experimentation. ^86^Rb^+^ efflux: The experiments were initiated by placing the choroid plexus in HEPES-aCSF containing 1 μCi/ml ^86^Rb^+^ and 4 μCi/ml ^3^H-mannitol at 37°C for 10 min to allow isotope accumulation in the tissue. The choroid plexus was briefly washed (2 × 5 s) in 37°C HEPES-aCSF prior to transfer (at 10 s intervals) to different HEPES-aCSF solutions (37°C) containing 20 μM bumetanide (B3023, Sigma) or DMSO vehicle (0.05%) for a total of 40 s, choroid plexuses from each animal randomly allocated to each condition. The efflux medium from each of the solutions was transferred into separate scintillation vials, as was the choroid plexus, the latter dissolved in 100 μl Solvable (6NE9100, Perkin Elmer). The ^86^Rb^+^ counts were corrected for ^3^H-mannitol counts (extracellular background), and the natural logarithm of the choroid plexus content A_t_/A_0_ was plotted against time [19] to obtain the ^86^Rb^+^ efflux rate (min^-1^) by linear regression analysis.

For the light and dark phase efflux experiment 12 animals was used (six either phase), but one dark phase lateral choroid plexus (bumetanide) was lost during experimentation. 500 μl of Ultima Gold™ XR scintillation liquid (6012119, PerkinElmer) was added to all scintillation vials and the radioactive content quantified in a Tri-Carb 2900TR Liquid Scintillation Analyzer (Packard).

### Statistics

Data were analysed with GraphPad Prism 9.0 (GraphPad Software, San Diego, US). All data are shown as mean ± SD and P < 0.05 was considered statistically significant. All data were tested for normal distribution using the Shapiro-Wilk test and data sets were tested for outliers (Grubbs’ test) when required, as indicated in figure legends. For normally distributed data, a one-sided or a two-tailed t-test was conducted, otherwise either a Mann-Whitney (unpaired), or Wilcoxon (signed rank) (paired) test. RNAseq was evaluated with the Welch test. If more than two parameters were compared (isotope experiments), a one-way ANOVA was employed. Correlation analyses were performed with Pearson’s test (telemetric data) or Spearman’s test (patient data). All statistical tests are indicated in the figure legends.

## RESULTS

### Dark phase increase in human ICP

Determination of ICP relies on surgical placement of a pressure probe and hence cannot be readily conducted in healthy individuals. To that end, we enrolled patients admitted to the Neurointensive Unit at Copenhagen University Hospital – Rigshospitalet, Copenhagen, Denmark with conditions (see *Methods*) that all required continuous monitoring of their ICP, as well as insertion of an external ventricular drain (EVD) to relieve the patient from ICP increases by draining CSF. The ICP was measured during at least four days of unchanged EVD counter-pressure (to avoid external sources of ICP changes) and initially averaged over the entire monitoring period in individual patients (7.67 ± 4.50 mmHg, n = 24). Separation of the ICP measurements within the light phase (6 AM to 6 PM) and the dark phase (6 PM to 6 AM) across the whole patient group revealed a higher ICP in the dark phase (8.51 ± 4.57 mmHg vs 6.83 ± 4.27 mmHg in the light phase, n = 24, P < 0.001, Fig. 1A, left panel). Determination of the mmHg change in ICP in the dark phase in the individual patient revealed a 30.0 ± 7.7% increase at night corresponding to an average nightly ICP elevation of 1.68 ± 0.48 mmHg compared to daytime (n = 24, P < 0.001, Fig. 1A, right panel). To obtain a relation to the patient blood pressure, we quantified this physiological parameter within both phases and observed an increase in average mean arterial pressure during the dark phase (90.7 ± 8.0 mmHg vs 88.3 ± 7.2 mmHg in the light phase, n = 21, P < 0.01, Fig. 1B, left panel) with an individual patient increase of 2.71% ± 0.73% in the dark phase (with an average nightly blood pressure elevation of 2.39 ± 0.66 mmHg (n = 21, P < 0.01, Fig. 1B, right panel). To determine if the blood pressure increase at night could underlie the nightly ICP increase, assuming that autoregulation was impaired, we performed correlation analysis on these paired data sets in each individual patient (Supplementary Fig. 1A) but observed no significant correlation between these parameters (n = 21, r^2^ = 0.05, P = 0.33).

**Figure 1.**
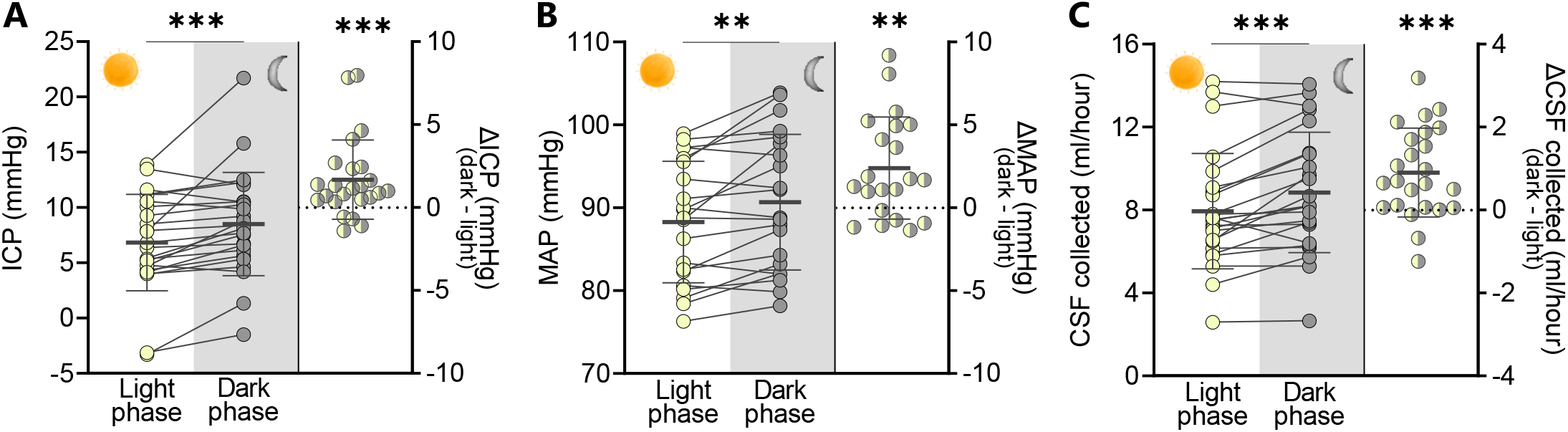
Dark-light phase fluctuations in patient ICP, blood pressure, and CSF dynamics in patients. **A**. Average ICP measured in patients during the light phase (6 AM to 6 PM) and the dark phase (6 PM to 6 AM), left panel, with the intra-patient ΔICP illustrated in the right panel, n = 24. Statistical significance evaluated with paired Wilcoxon t-test (left panel) and Wilcoxon signed-rank test (right panel). **B**. MAP measured in the same patient group obtained in the light phase (6 AM to 6 PM) and the dark phase (6 PM to 6 AM), left panel, with the intra-patient ΔMAP illustrated in the right panel, n = 21. Statistical significance evaluated with paired two-way t-test (left panel) and one sample t-test (right panel). **C**. CSF volume collected from the extraventricular drain in the patient group during the light phase (6 AM to 6 PM) and the dark phase (6 PM to 6 AM), left panel, with the intra-patient ΔCSF volume illustrated in the right panel, n = 24. Statistical significance evaluated with paired two-way t-test (left panel) and one sample t-test (right panel). ICP = intracranial pressure, MAP = mean arterial pressure, CSF = cerebrospinal fluid. Data are shown as mean ± SD. **P < 0.01, ***P < 0.001.

### Dark phase increase in human CSF dynamics

To determine if the nightly elevation in ICP could be a functional consequence of altered CSF dynamics in the dark phase, we took advantage of the CSF collection via the EVD as a proxy read-out of CSF secretion. To ensure stable measurements for each patient, we included only patients with consecutive light and dark measurements with the same EVD resistance (≥ four days with ≤ 10% difference in daily CSF secretion, see *Methods*). The CSF collection volume was higher in the dark phase (8.85 ± 2.85 ml/hour vs 7.94 ± 2.89 ml/hour in the light phase, n = 24, P < 0.001, Fig. 1C, left panel). Determination of the increase in CSF flow in the dark phase in the individual patient revealed a 12.4 ± 2.9% fluctuation with an average nightly elevation in CSF collection of 0.90 ± 0.21 ml/hour (n = 24, P < 0.001, Fig. 1C, right panel). Our data indicate that a nightly increase in CSF secretion rate could contribute to the elevation of ICP at night.

### Dark phase increase in rat ICP

To determine if the observed changes in human ICP and CSF flow within the dark/light phases could be observed in an animal model, which would allow subsequent delineation of the underlying molecular mechanisms, we employed rat experiments. ICP recordings were obtained in freely moving, non-anaesthetized rats with dual telemetric recordings from pressure probes placed on the epidural surface of the brain (ICP) and in the aorta (mean arterial pressure, MAP [19]). The average 24 h ICP of the experimental rats at the onset of the experimental window was 3.91 ± 1.13 mmHg, n = 9. Continuous ICP measurements across four days demonstrated clear diurnal fluctuations with a gradual increase from the initiation of dark phase and an abrupt decline with light onset (Fig. 2A). The ICP obtained in the dark phase was higher than in the light phase (4.34 ± 1.08 mmHg in the dark phase vs 3.48 ± 1.00 mmHg in the light phase, n = 9, P < 0.001, Fig. 2B, left panel). Determination of the ICP increase during the dark phase in the individual rat revealed an average ICP fluctuation of 27.1 ± 3.6% with a nightly elevation of 0.86 ± 0.05 mmHg (n = 9, P < 0.001, Fig. 2B, right panel). Analysis of the ICP bins at 8-10 hours after dark phase initiation (ZT 20-22 h; 4.59 ± 1.07 mmHg) and light phase initiation (ZT 8-10 h; 3.53 ± 0.99 mmHg, n = 9, P < 0.001, Fig. 2C, left panel) revealed an increase in ICP in the dark phase bin, with a difference in the individual rats of 33.3 ± 5.7% (1.05 ± 0.09 mmHg, n = 9, P < 0.001, Fig. 2C, right panel). The ICP thus increases in the dark phase to a similar extent in humans and rodents.

**Figure 2.**
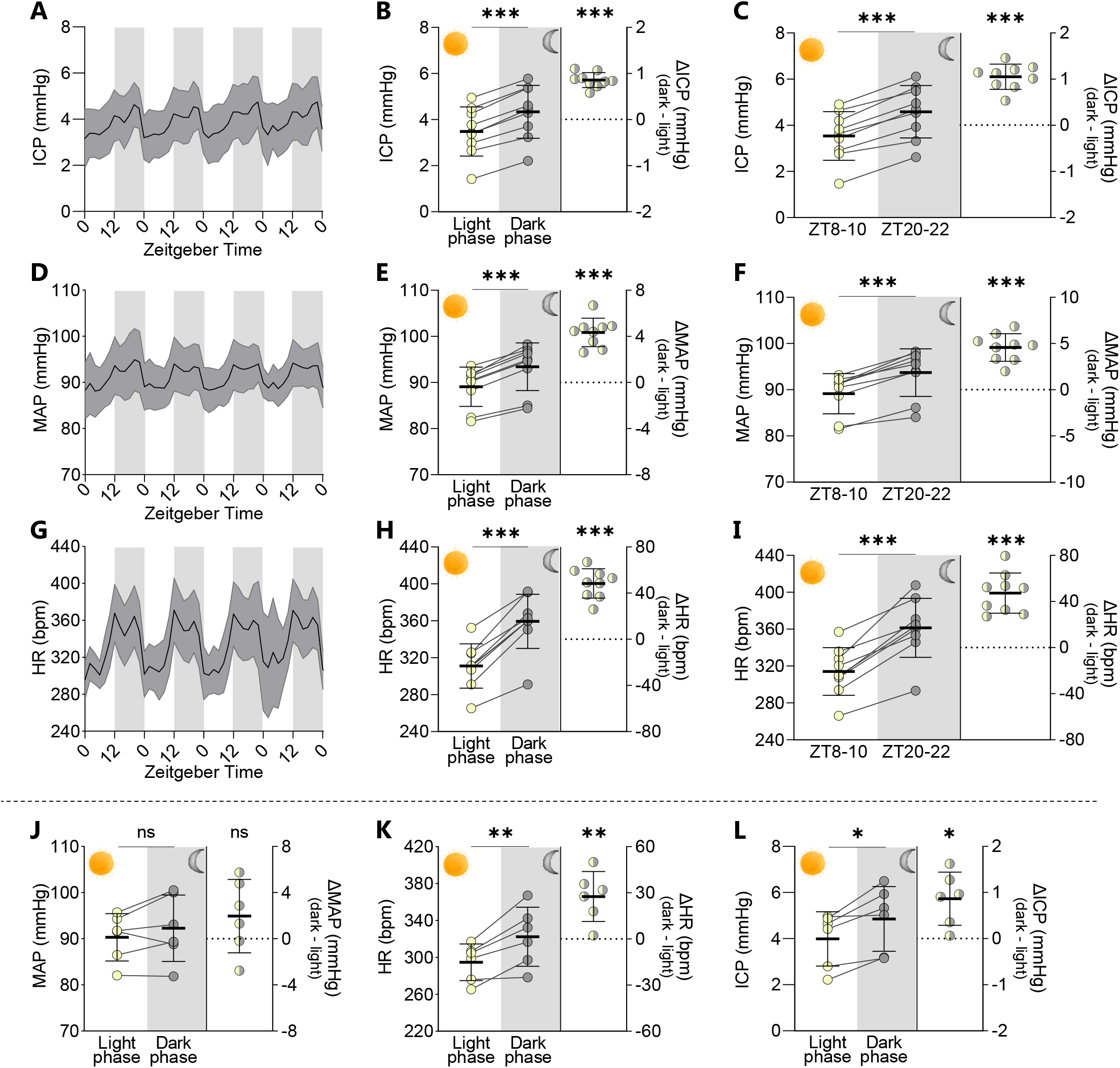
Dark-light phase rodent fluctuations in ICP, blood pressure and heart rate in rats. **A**. Continuous telemetric ICP recordings from freely moving, non-anaesthetized rats obtained in the light phase (ZT 0-12 h) and the dark phase (ZT 12-24 h), n = 9. **B** Average ICP across the light and dark phases illustrated in the left panel, with the intra-rat Δ ICP illustrated in the right panel, n = 9. **C**. The ICP at ZT 8-10 h and ZT 20-22 h (left panel) and the intra-rat Δ ICP (right panel), n = 9. **D**. Continuous telemetric MAP recordings from freely moving, non-anaesthetized rats obtained in the light phase (ZT 0-12 h) and the dark phase (ZT 12-24 h), n = 9. **E**. Average MAP across the light and dark phases illustrated in the left panel, with the intra-rat Δ MAP illustrated in the right panel, n = 9. **F**. The MAP at ZT 8-10 h and ZT 20-22 h (left panel) and the intra-rat Δ MAP (right panel), n = 9. **G**. Continuous telemetric heart rate recordings from freely moving, non-anaesthetized rats obtained in the light phase (ZT 0-12 h) and the dark phase (ZT 12-24 h), n = 9. **H**. Average heart rate across the light and dark phases illustrated in the left panel, with the intra-rat Δ heart rate illustrated in the right panel, n = 9. **I**. The heart rate at ZT 8-10 h and ZT 20-22 h (left panel) and the intra-rat Δ heart rate (right panel), n = 9. **J**. MAP, heart rate (**K**), and ICP (**L**) obtained in video-based identified time points of voluntary sedentary periods in windows spanning ZT 4-11 h in the light phase and ZT 16-23 h in the dark phase. Measurements in left panels and intra-rat delta values in right panels, n = 6. ICP = intracranial pressure, MAP = mean arterial pressure, HR = heart rate, ZT = zeitgeber time. Data are shown as mean ± SD and statistical significance was evaluated with paired two-tailed t-test (left panels) and one sample t-test (right panels). *** P < 0.001, ** P < 0.01, *P < 0.05, ns; not significant.

### Dark phase increase in rat ICP independently of activity level

With rats being predominantly nocturnal creatures [27], their elevated ICP could potentially arise from increased physical activity in this phase. To monitor their physical activity, we took advantage of the vascular pressure probe to concomitantly monitor the mean arterial pressure and heart rate in the experimental rats. These cardiovascular parameters essentially mimicked those of the ICP with stable diurnal fluctuations demonstrating elevated values during the dark phase (Fig. 2D and G). The mean arterial pressure obtained in the dark phase was 5% higher than in the light phase (93.4 ± 4.8 mmHg in the dark phase vs 89.1 ± 4.0 mmHg in the light phase, n = 9, P < 0.001, Fig. 2E left panel), and the heart rate was 16% higher than in the light phase (359 ± 28 bpm in the dark phase vs 311 ± 23 bpm in the light phase, n = 9, P < 0.001, Fig. 2H, left panel). The mean arterial pressure obtained 8-10 hours after dark phase initiation (ZT 20-22 h; 93.7 ± 4.9 mmHg) was 5% higher than that obtained 8-10 hours after light phase initiation (ZT 8-10 h; 89.1 ± 4.1 mmHg, n = 9, P < 0.001, Fig. 2F left panel), with a 5.10 ± 0.51% difference in the individual rats with a dark phase increase of 4.57 ± 0.47 mmHg, n = 9, P < 0.0001, Fig 2F, right panel). The heart rate followed a similar pattern with the values obtained 8-10 hours after dark phase initiation (ZT 20-22 h; 361 ± 30 bpm) being 15% higher than those obtained 8-10 hours after light phase initiation (ZT 8-10 h; 314 ± 24 bpm, n = 9, P < 0.001, Fig. 2I, left panel), with a 15.1 ± 1.7% difference in the individual rats with a dark phase increase of 47.3 ± 5.5 bpm (n = 9, P < 0.001, Fig 2I, right panel). Such dark phase-related increases in cardiovascular parameters may associate with altered body posture during cage roaming in the rat active phase, which is likely to affect ICP [28–30]. To obtain a scenario with a specific time point at which to quantify and correlate these physiological parameters, six rats were filmed during the peak dark and light phases (ZT 16-23 and 4-11 h) and distinct windows at which the rats were voluntarily sedentary subsequently selected for analysis. In these brief intervals, the mean arterial pressure was not significantly higher during the dark phase (92.3 ± 6.6 mmHg) compared to the light phase (90.3 ± 4.7 mmHg, n = 6, P = 0.19, Fig. 2J), whereas the heart rate remained elevated (322 ± 29 bpm in dark phase vs 295 ± 18 bpm in light phase, n = 6, P < 0.01, Fig. 2K) and so did the ICP (4.85 ± 1.28 mmHg in the dark phase vs 3.99 ± 1.07 mmHg in the light phase, n = 6, P < 0.01, Fig. 2L). Notably, the dark phase increase in all parameters for the individual rats were smaller in the sedentary periods compared to the increase in the two-hour bins during the peak within the phase. Correlation analysis demonstrated no interaction between the nightly increase in mean arterial pressure and the increase in ICP, whether the animals were sedentary in the dark phase (n = 6, r^2^ = 0.06, P = 0.65) or not (n = 9, r^2^ = 0.20, P = 0.23, Suppl. Fig 1B). Taken together, these results suggest that the increased ICP observed in the rats during the dark phase resembles that of the human patients and is a constant feature that is not dictated by nocturnal increase in activity level or related fluctuations in cardiovascular parameters.

### Dark phase increase in rat CSF secretion rate

To determine whether an increased CSF secretion rate could underlie the elevated ICP in the dark phase, we determined the CSF secretion rate in anesthetized, thermo-controlled, and mechanically ventilated rats in the light phase (ZT 8 h) and dark phase (ZT 20 h), with the rats completely shielded from visible light both prior to the anesthesia induction and during the experimental procedure (see *Methods*). The CSF secretion rate was determined with ventriculo-cisternal perfusion where artificial CSF containing a fluorescent dye is infused into the lateral ventricle at a constant rate and CSF simultaneously sampled from a cisterna magna puncture. The CSF secretion rate can thus be obtained based on the dilution of fluorescent dye, which arises with endogenously formed CSF. Fig. 3A illustrates the fluorescent dilution in representative experiments conducted in the dark phase and in the light phase. The values obtained during the last 20 min of the experimental window were employed for quantification and are averaged in Fig. 3B. CSF secretion increased by 20% in the dark phase (5.82 ± 0.62 μl/min vs 4.84 ± 0.45 μl/min in the light phase, n = 5 in each phase, P < 0.05). The CSF secretion is thus, as also observed in the patient cohort, elevated during the dark phase and may contribute to the elevated ICP observed in both humans and rats.

**Figure 3.**
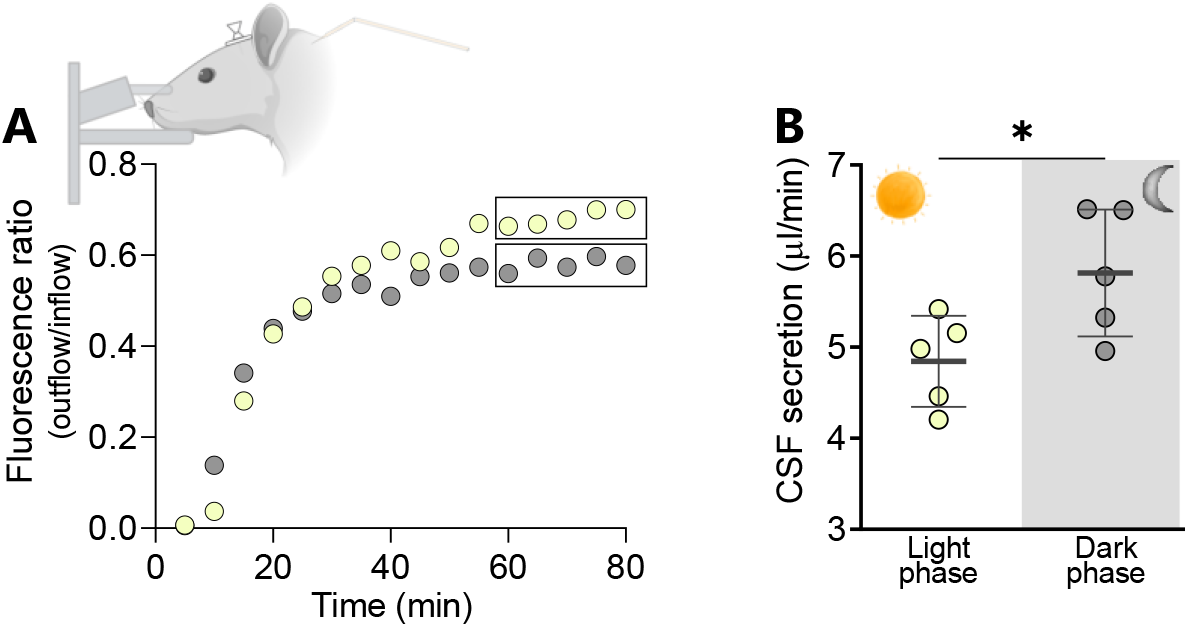
Dark-light phase fluctuation in rat CSF secretion rate. A. Representative time course of the dextran dilution quantified from the cisterna magna fluid collection taking place during the ventriculo-cisternal perfusion assay in mechanically-ventilated anaesthetized rats in the light phase (ZT 8 h, yellow spheres) and dark phase (ZT 20 h, grey spheres). Squares indicate the time interval employed for quantification of the CSF secretion rate illustrated in B, n = 5 of each. Data are shown as mean ± SD and statistical significance was evaluated with paired two-tailed t-test. *P < 0.05.

### Minor transcriptomic changes in the choroid plexus transporters involved in CSF secretion

To determine whether dark phase-modulation of the molecular apparatus supporting CSF secretion across the choroid plexus could underlie the elevated CSF secretion, we performed RNAseq of rat choroid plexus acutely isolated from rats in the light phase (ZT 8 h) and the dark phase (ZT 20 h). Quantification of transcripts coding for transporters involved in CSF secretion [31] (Fig. 4A), illustrates that two of these choroid plexus transporters were upregulated in the dark phase - the Na^+^-driven Cl^-^/HCO_3_ ^-^ exchanger NCBE (11%; 318 ± 19 TPM vs 287 ± 10 TPM in the light phase, n = 6, P < 0.05) and the electroneutral Na^+^,HCO_3_^-^ cotransporter 1 NBCn1 (21%; 8.69 ± 0.84 TPM vs 7.18 ± 0.53 TPM in the light phase, n = 6, P < 0.05, Fig. 4B), the latter of which is, however, expressed at a very low level and may not participate in CSF secretion [24]. Two transport mechanisms were even downregulated in the dark phase (aquaporin 1, 22 %; 472 ± 44 TPM vs 604 ± 38 TPM in the light phase, n = 6, P < 0.01) and the electrogenic Na^+^,HCO_3_^-^ cotransporter 2 NBCe2 (15% 540 ± 35 TPM vs 635 ± 15 TPM in the light phase, n = 6, P < 0.01, Fig. 4B), while some of these transcript levels displayed no sign of daily fluctuations (the Na^+^/K^+^-ATPase α1 subunit, the Na^+^,K^+^,2Cl^-^ cotransporter NKCC1, and the anion exchanger AE2, Fig. 4B). The stable expression of NKCC1 and the Na^+^/K^+^-ATPase in the light and dark phase was mirrored at the protein level, where there was no significant difference between protein expression as evaluated by western blotting of these two choroid plexus proteins on choroid plexus excised in the light phase (ZT 8 h) and the dark phase (ZT 20 h), n = 5-6 (Suppl. Fig. 2). Notably, regulatory properties of the choroid plexus CSF secretion machinery, such as the transient receptor potential vanilloid 4 (TRPV4) channel and the with-no-lysine kinase 1 (WNK1), modulate NKCC1 activity in choroid plexus via the Ste20/SPS1-related proline alanine-rich kinase (SPAK [32,33]). These two modulators were both upregulated approximately 30% in the dark phase (TRPV4; 79.6 ± 4.9 TPM vs 59.8 ± 5.4 TPM in the light phase, n = 6, P < 0.001 and WNK1 22.9 ± 0.9 TPM vs 17.3 ± 1.0 TPM in the light phase, n = 6, P < 0.001) and could, as such, contribute to the increase in CSF secretion during the dark phase.

**Figure 4.**
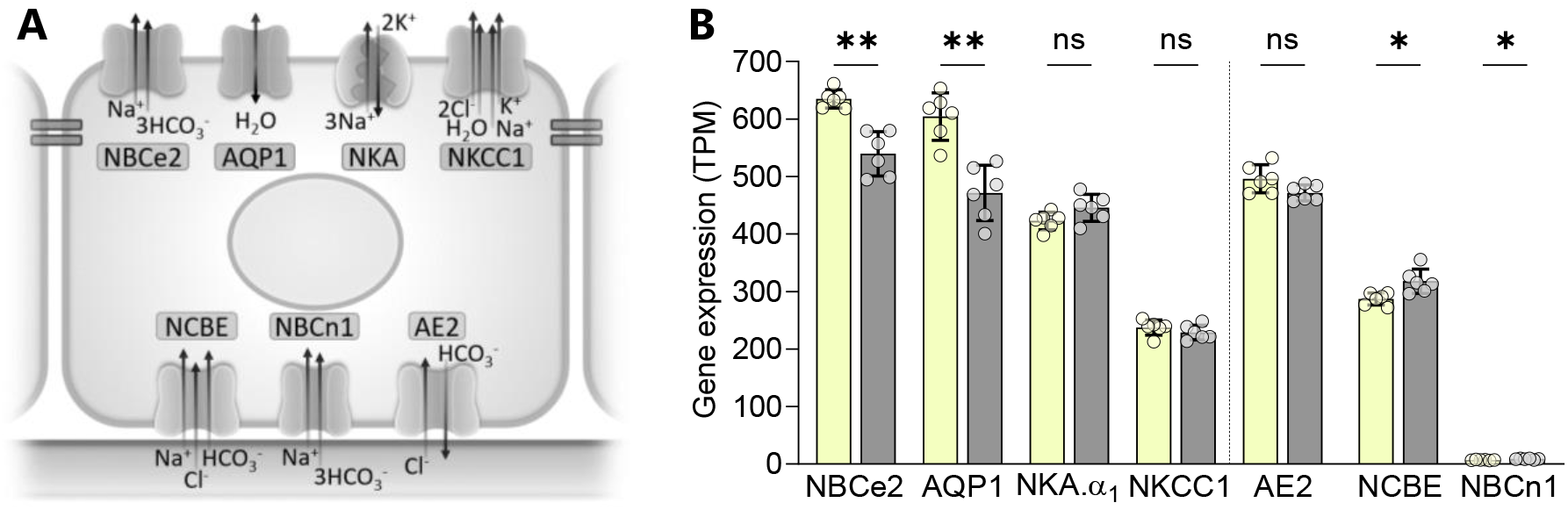
Dark-light phase fluctuation in rat choroid plexus transporter expression. **A. A** schematic choroid plexus epithelial cells with transporters proposed involved in CSF secretion illustrated on the respective membranes. **B**. Gene expression of these transporters quantified as transcripts per million (TPM) following RNAseq of acutely excised choroid plexus tissue obtained in the light phase (ZT 8 h, yellow bars) and the dark phase (ZT 20 h, grey bars), n = 6 of each. Transcripts left of the dashed vertical line are those of transporters expressed in the luminal membrane. NBCe2 = the electrogenic Na^+^,HCO_3_^-^ cotransporter 2, AQP1 = aquaporin 1, NKA = the Na^+^/K^+^-ATPase.α_1_ subunit, NKCC1 = the Na^+^, K^+^, 2Cl^-^ cotransporter, NCBE = the Na^+^-driven Cl^-^/HCO_3_^-^ exchanger, NBCn1 = the electroneutral Na^+^,HCO_3_^-^ cotransporter 1, and AE2 = the anion exchanger AE2, ZT = zeitgeber time. Data are shown as mean ± SD and statistical significance was evaluated with Welch ANOVA test. ** P < 0.01, *P < 0.05, ns; not significant.

### Choroid plexus transporter activity increased in the dark phase

With the dark phase transcriptional upregulation of modulating properties of the transport mechanisms in choroid plexus, the activity of choroid plexus transport mechanisms could be modulated in the dark phase despite their stable expression level. To determine whether two of the key components in CSF secretion, the Na^+^/K^+^-ATPase and the NKCC1 [27,28] display varied activity in choroid plexus during the light/dark phase, we performed isotope flux assays on choroid plexus tissue acutely excised from rats 8 hours after initiation of the light phase (ZT 8 h) and 8 hours after initiation of the dark phase (ZT 20 h), with care taken to protect the dark phase animals from light exposure prior to tissue isolation (see *Methods*). The Na^+^/K^+^-ATPase activity was determined with an isotope uptake assay based on the K^+^ congener ^86^Rb^+^, which readily replaces K^+^ in the transport cycle (Fig. 5A). The total ^86^Rb^+^ uptake in the dark phase was 1.54 ± 0.45 × 10^4^ cpm in control solution vs 0.26 ± 0.05 × 10^4^ cpm in the presence of the Na^+^/K^+^-ATPase inhibitor ouabain (n = -6, P < 0.001, Fig. 5B) and in the light phase it was 2.02 ± 0.44 × 10^4^ cpm in control solution vs 0.28 ± 0.02 × 10^4^ cpm in the presence of ouabain (n = 6, P < 0.001, Fig. 5B). The Na^+^/K^+^-ATPase-mediated activity was obtained as the ouabain-sensitive fraction of the uptake, which was not significantly different between the dark and the light phase (1.40 ± 0.35 × 10^4^ cpm in the dark phase vs 1.75 ± 0.45 ×10^4^ cpm in the light phase, n = 5-6, P = 0.86, Fig. 5C). The NKCC1 activity was determined with an isotope efflux assay of ^86^Rb^+^ pre-equilibrated in the excised choroid plexus (Fig. 5D). The total ^86^Rb^+^ efflux in the dark phase was 0.52 ± 0.04 min^-1^ in control solution vs 0.20 ± 0.01 min^-1^ in the presence of the NKCC1 inhibitor bumetanide (n = 5-6, P < 0.001, Fig. 5E-F) and in the light phase it was 0.45 ± 0.06 min^-1^ in control solution vs 0.21 ± 0.02 min^-1^ in the presence of bumetanide, (n = 5-6, P < 0.001, Fig. 5E-F). The bumetanide-sensitive fraction of the ^86^Rb^+^ uptake reflects the NKCC1-mediated transport activity, which was elevated 44% in the dark phase (0.33 ± 0.05 min^-1^ vs the light phase 0.23 ± 0.06 min^-1^, n = 5, P = < 0.05, Fig. 5G). These data indicate that the elevated rate of CSF secretion during the dark phase may, in part, be a result of elevated activity of the choroid plexus cotransporter NKCC1.

**Figure 5.**
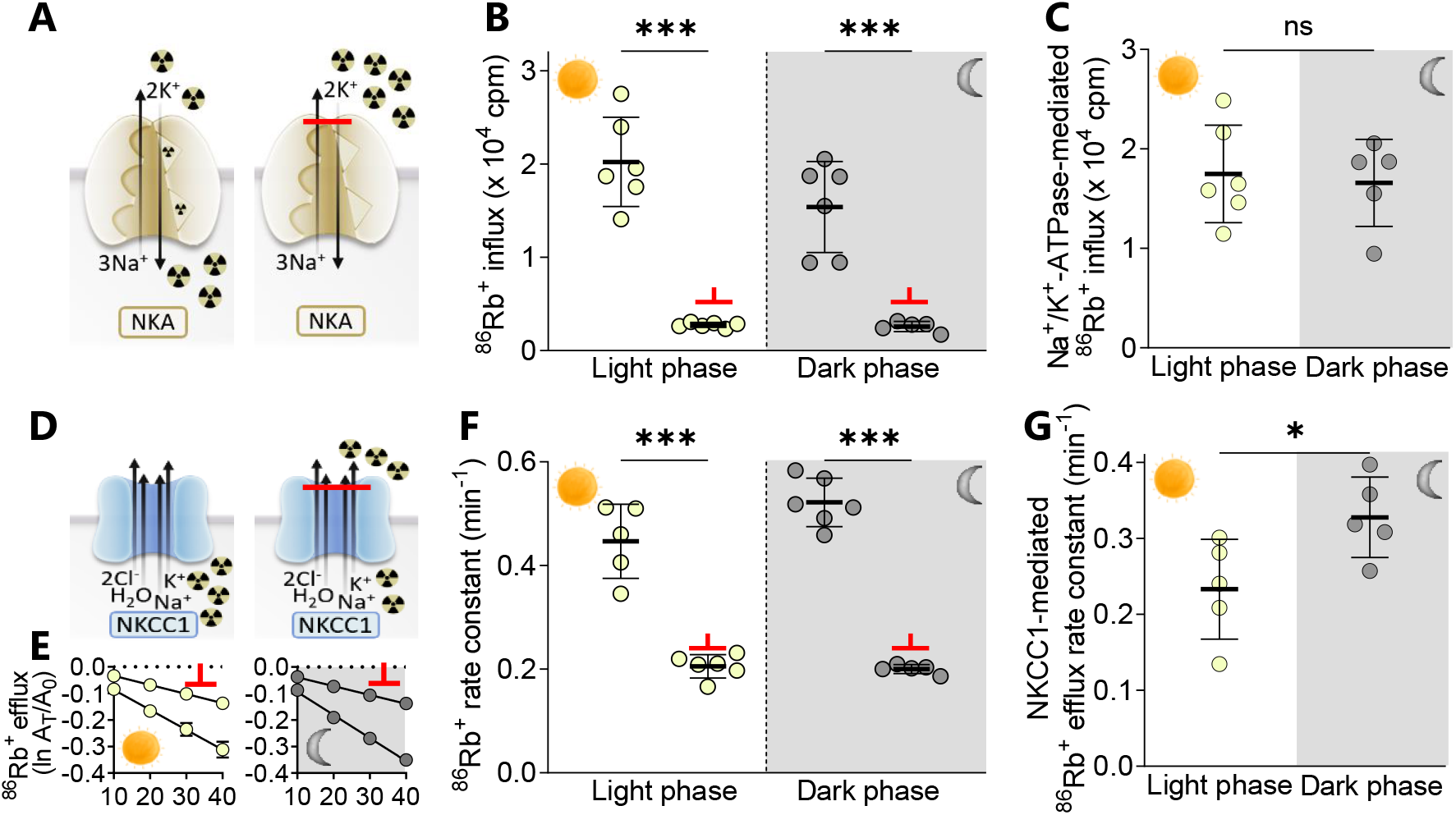
Dark-light phase fluctuation in rat choroid plexus transporter function. **A**. Schematic illustrating uptake of ^86^Rb^+^ via the Na^+^/K^+^-ATPase (left panel), inhibited by ouabain (2 mM, right panel). **B**. ^86^Rb^+^ uptake in choroid plexus excised during the light (ZT 8 h) and dark phase (ZT 20 h) in the absence and presence of ouabain (marked with a red bar), n = 5-6. **C**. The Na^+^/K^+^-ATPase-mediated ^86^Rb^+^ influx was obtained from the difference between control and ouabain-treated choroid plexus within the same rat, n = 5-6. **D**.Schematic illustrating efflux of ^86^Rb^+^ via NKCC1 (left panel) and inhibited by bumetanide (20 μM, right panel). **E**. ^86^Rb^+^ efflux from choroid plexus excised during the light (ZT 8 h) and dark phase (ZT 20 h) in the absence and presence of bumetanide (marked with a red bar), n = 5-6, one outlier removed from the light phase control group. The Y-axis represents the natural logarithm of the ^86^Rb^+^ amount left in the choroid plexus at time T (A_T_) divided by the initial amount at time 0 (A_0_). **F**. Efflux rate constants calculated for the data displayed in E. **G**. The NKCC1-mediated ^86^Rb^+^ efflux was obtained from the difference between control and bumetanide-treated choroid plexus within the same rat, n = 5. NKA = the Na^+^/K^+^-ATPase, NKCC1 = the Na^+^, K^+^, 2Cl^-^ cotransporter. Data are shown as mean ± SD and statistical significance was evaluated with one-way ANOVA followed by Tukey’s multiple comparison test (B and F) or unpaired t-test (C and G). *** P < 0.001, *P < 0.05, ns; not significant.

## DISCUSSION

Here, we demonstrate that the dark phase-increase in ICP is associated with elevated CSF secretion in both humans and rats, irrespective of their diurnal/nocturnal activity cycle, in part due to elevated transport activity of the choroid plexus. The ICP has long been thought to increase at night in humans [4–6]. However, most studies associate with the caveat that the human subjects undergoing ICP measurements, as part of their diagnostic work up or treatment, change posture when switching to horizontal position during sleep. Such transition will artificially increase the nightly ICP measurements, as ICP increases with tilting in a sense that the more horizontal the subject, the higher the ICP, and even more pronounced in positions with the head further lowered [7,8]. We here took advantage of a patient group in the neuro intensive care unit, who were largely bedridden due to the severity of their disorders, and thus residing in (near)horizontal positions during the data acquisition. The patient ICP was continuously monitored for diagnostic purposes and revealed a 30% increase at night. The patient blood pressure displayed a slight, but statistically significant parallel nightly increase (3%). Such dark phase increase may usually be masked by altered body position and associated elevated activity in the daytime. However, we observed no correlation between the nightly increase in these two pressure parameters (ICP and MAP), suggesting that a potential diurnal regulation of blood pressure is not driving the nightly elevation of ICP. The data align with earlier demonstrations that ICP does not increase with elevated blood pressure [34,35]. Freely moving experimental rats equipped with telemetric probes displayed a mirror increase in dark phase ICP (30%) and blood pressure (5%), the former fluctuation in agreement with visible inspection of previously published continuous telemetric data sets [19,28]. Although the rodent measurements have the advantage of being recorded on quadruped, thus mostly horizontal, animals with lesser body position-confounding elements, the nocturnal rodent nature prompts an elevated activity level during the dark phase. Such elevated activity could potentially modulate the ICP, albeit in both directions: an increased activity could include i) raising the torso/head, which lowers the rat ICP [28–30] versus ii) elevated blood pressure, which could potentially increase the rat ICP. However, analogous to the human data, the dark phase increase in these two pressure parameters (ICP and MAP) did not display correlation in the rats, suggesting that a blood pressure increase, potentially arisen with the increased nocturnal activity, did not drive the elevated ICP. In support thereof, in windows of voluntary sedentary periods in the otherwise active dark phase, the ICP remained elevated. Taken together, the ICP increases in the dark phase in rats and humans, apparently independently of body posture and diurnal/nocturnal activity preferences and, in addition, not originating from dark/light-mediated changes in systemic arterial blood pressure.

With our previously published proof of concept that modulation of the CSF secretion rate in rodents directly affect ICP [19,36], we hypothesized that a dark phase increase in CSF dynamics could underlie the elevated ICP. Here, we took advantage of the included patient group being equipped with external ventricular drains to allow continuous CSF collection to clinically manage the ICP within a tolerable range. The CSF volume collected was determined at 12-hour intervals (night/day) and represents a proxy for CSF secretion in that interval. Of note, the collected volume does not represent the complete CSF secreted, as the usual drainage pathways are still, at least to some extent, functional. The 200 ml/day CSF collected therefore is much reduced compared to the approximately 500 ml/day secreted in the adult human [37]. Nevertheless, the CSF volume collected at night was 12% higher than the day time volume in humans, which aligns with a previous MRI-based study on healthy volunteers that demonstrated nightly increases in net CSF flow through the aqueduct of Sylvius [12]. A subsequent study employing non-triggered phase-contrast MRI failed to reproduce this nightly fluctuation [13]. It is noteworthy that the time window including the peak CSF flow (12-4 am) in the former study [12] was not tested in the latter[13], which may then have missed the peak period. Our indirect measurement of CSF secretion could also be affected by day-night differences in CSF drainage capacity [38,39], and we therefore determined the CSF secretion in anaesthetized rats in the peak of each phase (light/dark). Great efforts were made to ensure that the rats were exposed to minimal light during the dark phase experiments (see *Methods*). Analogous to the human data, the CSF secretion rate was higher (20%) in the dark phase than in the light phase, suggesting that an elevated rate of CSF secretion in the dark phase, irrespective of diurnal/nocturnal activity, could be a direct contributing factor to the increased ICP in the dark phase. The CSF secretion rate was determined in the anesthetized rats with the ventriculo-cisternal perfusion assay, which relies on intraventricular infusion of dextran-containing artificial CSF at a fixed rate, with concomitant CSF collection from a cisterna magna puncture. The dextran is diluted by de novo CSF secretion, the rate of which can thus be determined from its dilution factor through the ventricular system [20,21]. Some researchers have expressed concern about this technique [40–42], one of which based on an assumption that the dextran enters the brain parenchyma during the experimental procedure, which would cause an artificially low amount of dextran to exit the cisterna magna puncture. We cannot detect any dextran in the brain parenchyma of rats following execution of a full experimental procedure [36] and suspect that Liu et al. [43] detected such parenchymal tracer entry due to the lack of a cisterna magna puncture in their experimental approach. Their constant tracer infusion in a mouse lateral ventricle (42 μl in total) with no experimentally-induced exit path for the excess brain fluid could well promote their observed fluorescent entry into the brain tissue. Secondly, the proposed alteration in ventricular osmolarity and temperature with the infused artificial CSF [41] is circumvented in our experiments with careful matching of the CSF osmolarity [36] and pre-heating of the gas-equilibrated HCO_3_^-^ -based CSF immediately prior to entry into the ventricles. However, with acknowledgement of the absolute quantification of CSF secretion rates in different animals remaining unsettled, our data simply address the dark/light phase *variation* in the CSF secretion rate, which is independent of the exact CSF secretion rate, and here demonstrated to increase at night.

The ventriculo-cisternal perfusion assay quantifies the CSF secretion irrespective of origin of the CSF. However, with the majority of the CSF secreted by the choroid plexus [38], which exhibits circadian fluctuation of known clock genes [44,45], we determined the dark/light phase transcriptomic regulation of the transport mechanisms known to be implicated in CSF secretion by the choroid plexus [31,37]. With no robust upregulation at the mRNA transcript level of any of these, we propose a dark phase increase in factors implicated in post-translational regulation of these transport mechanisms. We observed a 30% dark phase upregulation of TRPV4 and WNK1. The former increases CSF secretion by a direct activation of NKCC1, via the WNK/SPAK signaling pathway, which the latter feeds into [32,33]. We, accordingly, detected an increased transport activity of NKCC1, and not the Na^+^/K^+^-ATPase, in choroid plexus excised in the dark phase compared to that obtained in the light phase. We therefore propose that the dark phase elevation of ICP observed in humans and rats, at least in part, originate from an associated increase in the rate of CSF secretion, part of which is likely to occur with an elevated transport activity of the choroid plexus NKCC1. Curiously, the elevated CSF dynamics occurred in the dark phase in both species, irrespective of the human diurnal activity vs the rat nocturnal activity. This finding suggests that the CSF dynamics are governed by the circadian rhythm, rather than induced directly by the action of sleep and its associated cellular and physiological changes.

CSF drainage into the lymph nodes, and further to the vasculature, peaks during the dark (active) phase in anesthetized mice [39] and when mice transition from anesthesia to awake (in mimic of sleep-to-awake and thus light phase-to-dark phase [38]). The day/night drainage capacity cycle appears to be governed by the circadian rhythm, rather than sleep [39], as also here observed for the CSF secretion capacity and ICP fluctuations. Elevated CSF drainage in the equivalent of the dark phase in these mice aligns with the elevated CSF secretion here observed in rats and humans in the dark phase. The matched CSF in- and outflow capacity is indicated in the constant whole brain water percentage across the light/dark phase in the rats (Suppl. Fig. 3). The total brain water may, however, distribute differently *within* the brain with the dark/light phases, as enhanced CSF drainage to the cervical lymph nodes appears to occur in antiphase to CSF entry into the brain tissue along the paravascular spaces of the penetrating arterioles [39].

Limitations to this study include employing a patient group rather than healthy control individuals, which was necessary to obtain access to ICP measurements and CSF collection. The patient group was characterized by acute brain injury; however, this term covers different pathophysiological processes depending on the underlying specific disease, although increased ICP is part of the final common pathway for all these diseases. The clinical treatment was generally aimed at controlling ICP, but treatment guidelines did not depend on the time of day; in fact, it is likely that clinical management may have ‘buffered’ natural changes in ICP. The collected CSF was used as a surrogate marker of the CSF secretion rate; however, although the drainage pathways may well have retained (part of) their capacity in the given setting, they may not depend on time of day. Thus, we believe that day-night fluctuations are well reflected by the collected volumes. Another limitation to the study is the lack of information regarding sleeping patterns and phases, which were measured neither in the patient cohort nor in rats. Regardless of sedation, sleep disturbances are prevalent and severe in generally critically ill patients [46,47] and probably also in patients with severe acute brain injury; our data thus suggest that normal sleep is not a requirement for day-night ICP fluctuations. This could not be assessed in the experimental rats, as these were anesthetized in both phases.

## CONCLUSIONS

In conclusion, our data demonstrate that CSF secretion is elevated in the dark phase in humans and rodents, in part due to elevated activity of a choroid plexus transporter. The altered CSF flow is suggested to serve as a mediator of the detected dark-phase ICP elevation in both species, *irrespective of nocturnal/diurnal activity preferences* and bipedal/quadrupedal posture. The elevated ICP therefore does not appear to associate with sleep, but rather dictated by the circadian rhythm. This retained circadian pattern may stem from the highly conserved choroid plexus originating from a common ancestor with a predicted diurnal activity preference [48]. With the reversibility of sleep rhythms, these may well have been modulated to fit with the environmental predation risk, as species evolved [49–51]. It is thus likely that the light/dark phase rhythm of the CSF secretion capacity of the choroid plexus has not adjusted with the nocturnal activity pattern observed in the rats. Future research is predicted to unravel the molecular pathways coupling the light/dark cycle to altered CSF dynamics and associated pressure. One may tap into such CSF-flow regulatory candidates with the ongoing aim to reveal specific and efficient pharmaceuticals targeted towards the CSF secretion apparatus in disorders involving elevated ICP.

## DECLARATIONS

### Ethics approval and consent to participate

Patient data were, retrospectively, acquired from the Neurointensive Unit at Copenhagen University Hospital – Rigshospitalet, Copenhagen, Denmark with approval by the Danish National Committee on Health Research Ethics (Approval No. R-21040902) and the Danish Data Protection Agency (P-2021-565). All animal experimental work conformed to the legislations for animal protection and care in the European Community Council Directive (2010/63/EU), and approved by the Danish Animal Experiments Inspectorate (license no. 2016-15-0201-00944 and 2018-15-0201-01595).

### Consent for publication

Not applicable

### Availability of data and materials

We confirm that all data from this study will be available upon request.

### Authors’ contributions

A.B.S. and N.M. designed the research; A.B.S., B.L.E, D.B., and S.N.A performed research, M.H.O and K.M. collected clinical data, A.B.S., B.L.E., D.B., M.H.O., AND S.N.A. analyzed the data; B.L.E. and N.M. drafted the manuscript. All authors contributed to the manuscript and approved the final version.

## Acknowledgements

We are grateful for the technical assistance from technician Trine L Devantier.

