## Supplemental figures for "Nocturnal increase in cerebrospinal fluid secretion as a regulator of intracranial pressure"

### SUPPLEMENTARY FIGURES

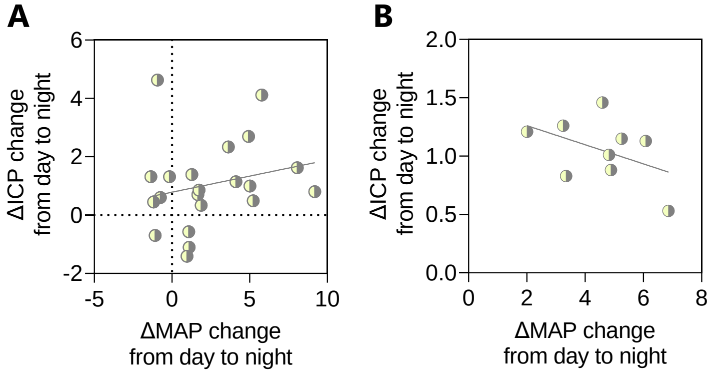


**Supplementary figure 1.** Correlation analysis of the nightly change in paired intracranial pressure (ICP) and mean arterial pressure (MAP) from **A,** patient cohort (n = 21, $r^2$ = 0.05, P = 0.33, Spearman correlation) and **B.** freely behaving rats (n = 6, $r^2$ = 0.06, P = 0.23, Pearson correlation).

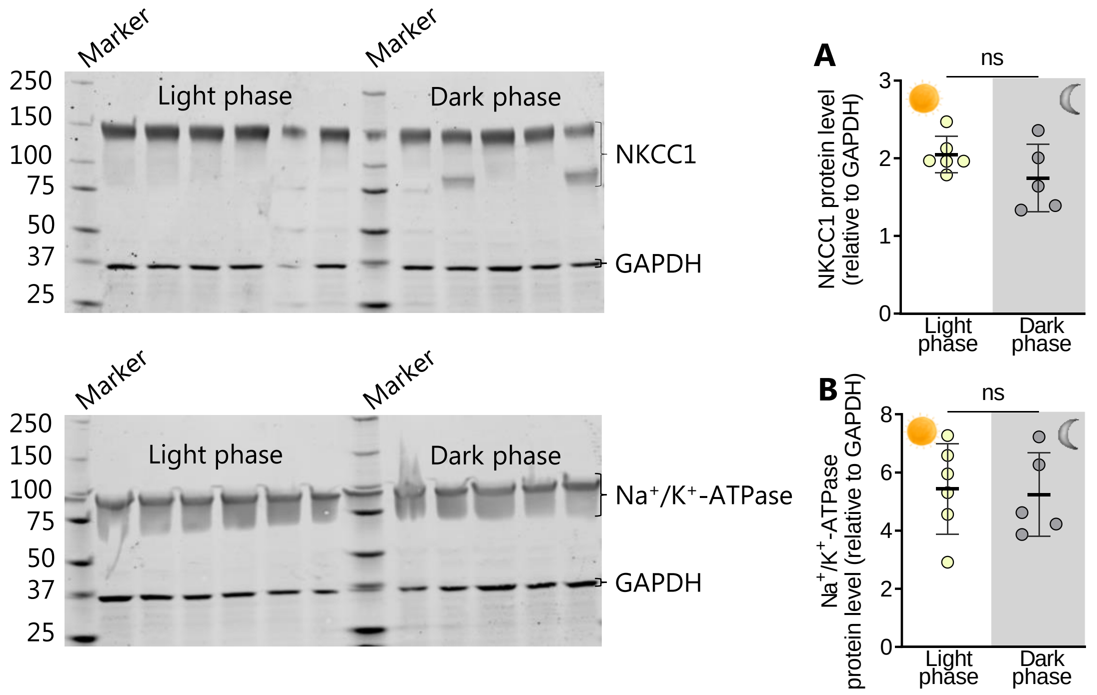


**Supplementary figure 2.** $Na^+$,$K^+$,$2Cl^-$ cotransporter (NKCC1), and $Na^+/K^+$-ATPase protein expression in choroid plexus acutely excised in the light and the dark phase. Analyzed by SDS-PAGE (Mini-protean TGX, precast 4-20%, Bio-Rad) and immunoblotting (Immobilon-FL membranes, Millipore, MA), left panel, utilizing glyceraldehyde 3-phosphate dehydrogenase (GAPDH) as a loading control. Primary antibodies included sheep anti-NKCC1, 1:400 (SO22D, Dundee, anti-sheep), mouse anti-$Na^+/K^+$-ATPase-$\alpha$1,1:60 (a6F, DSHB), chicken anti-GAPDH, 1:800 (AB2302, Millipore) and detection performed using fluorophore-conjugated secondary antibodies (LI-COR) and scanned on an Odyssey CLx imaging system. No significant difference was observed between the two phases for **A,** NKCC1 (dark phase, 1.7 ± 0.4, n = 5, vs light phase, 2.0 ± 0.2, n = 6, P = 0.18), or **B,** $Na^+/K^+$-ATPase α1 (dark phase, 5.2 ± 1.4, n = 5, vs light phase, 5.4 ± 1.6, n = 6, P = 0.84). Data are presented as mean ± SD, and statistical significance ($P < 0.05$) determined using unpaired t-test. ns; not significant.

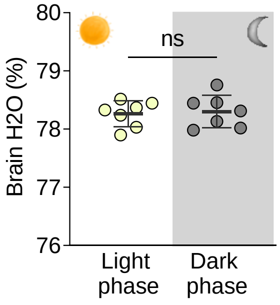


**Supplementary figure 3.** Quantification of brain water content in brains excised during the light (ZT 8h) and the dark (ZT 20h) phase. Swiftly excised rat brains were weighed, dried at 100°C for a minimum of 72 h before re-weighing. Brain water content in percentage did not differ between phases (dark phase 78.3 ± 0.3 % vs light phase 78.3 ± 0.2 %, $n = 7$, $P = 0.78$). Data presented as mean ± SD and statistical significance determined using unpaired t-test. ns; not significant.
